# Comprehensive analysis of human hookworm secreted proteins using a proteogenomic approach

**DOI:** 10.1101/406843

**Authors:** J Logan, SS Manda, YJ Choi, M Field, RM Eichenberger, J Mulvenna, SH Nagaraj, RT Fujiwara, P Gazzinelli-Guimaraes, L Bueno, V Mati, M Mitreva, J Sotillo, A Loukas

## Abstract

The human hookworm *Necator americanus* infects more than 400 million people worldwide, contributing substantially to the poverty in these regions. Adult stage *N. americanus* live in the small intestine of the human host where they inject excretory/secretory (ES) products into the mucosa. ES products have been characterized at the proteome level for a number of animal hookworm species, but until now, the difficulty in obtaining sufficient live *N. americanus* has been an obstacle in characterizing the secretome of this important human pathogen. Herein we describe the ES proteome of *N. americanus* and utilize this information to conduct the first proteogenomic analysis of a parasitic helminth, significantly improving the available genome and thereby generating a robust description of the parasite secretome. The genome annotation resulted in a a revised prediction of 3,425 fewer genes than initially reported, accompanied by a significant increase in the number of exons and introns, total gene length and the percentage of the genome covered by genes. Almost 200 ES proteins were identified by LC-MS/MS with SCP/TAPS proteins, ‘hypothetical’ proteins and proteases among the most abundant families. These proteins were compared to commonly used model species of human parasitic infections, including *Ancylostoma caninum, Nippostrongylus brasiliensis* and *Heligmosomoides polygyrus*. Our findings provide valuable information on important families of proteins with both known and unknown functions that could be instrumental in host-parasite interactions, including protein families that might be key for parasite survival in the onslaught of robust immune responses, as well as vaccine and drug targets.

## Introduction

Hookworm infection is one of the most pertinent and life-limiting parasitic infections worldwide, affecting more than 400 million people in tropical regions of Asia, Africa and South America (1, 2). Chronic infection with hookworms results in fatigue, abdominal pain, diarrhea, weight loss and anemia (3). In children, it can cause growth retardation and impairments in cognitive development (4), and in pregnant women these infections lead to poor birth outcomes including low birth weight, increased perinatal morbidity and mortality (5, 6). Moreover, hookworm infections result in 3.2 million disability-adjusted life years lost annually (7). Hookworm infection therefore contributes significantly to widespread poverty in the majority of endemic regions (8).

The life cycle of the most widespread of the anthropophilic hookworms, *Necator americanus*, is direct, with no intermediate hosts involved. Eggs are passed out of the body in the feces and, under favorable conditions, hatch releasing first-stage (L1). Larvae undergo several molts to reach the infective third (L3) stage which can penetrate human skin (9). Upon infection the parasite migrates through the circulatory system to the lungs, where it moves up the trachea and is eventually swallowed, thus commencing its final sojourn through the gastrointestinal tract to reside in the small intestine where mature adult worms can live for up to 10 years (9). The adult stage of *N. americanus* produces macromolecules known as excretory/secretory (ES) products, which consist of a battery of proteins that have evolved to interact with human host tissues and facilitate parasitism (10). These ES products have the potential to not only be targeted as potential vaccine and diagnostic candidates, but also to shed light on how these parasites evade immune destruction (11–14).

Despite their potential biotechnological utility, only a limited number of *N. americanus* ES proteins have been described to date, and most of them have been identified as cDNAs based on their homology to proteins from more readily accessible hookworm species from animals such as *Ancylostoma caninum* (15). In terms of vaccine antigens, a handful of *N. americanus* ES products including glutathione-S-transferases, aspartic proteases and sperm-coating proteins/Tpx-1/Ag5/PR-1/Sc7 (SCP/TAPS), have been identified at the cDNA level, and vaccine efficacy of recombinant proteins assessed in animal models and phase 1 clinical trials (13, 16, 17). SCP/TAPS, also referred to as venom allergen-like (VAL) or Activation-associated Secreted Proteins (ASPs) (Pfam accession number no. PF00188) have been reported from many helminths, but appear to be significantly expanded in the genomes and secreted proteomes of gut-resident clade IV and V parasitic nematodes, including the hookworms (11, 18–20).

Considerably little is known about the roles of *N. americanus* proteins in immunoregulation compared with many of the parasites that serve as animal models for human infections. Hsieh and colleagues reported an *N. americanus* protein(s) which selectively bound Natural Killer (NK) cells resulting in IL-2-and IL-12-dependent IFN-γ production, although the identity of the protein was not determined (21). Calreticulin, a protein from the ES products of *N. americanus*, was identified as having an immunomodulatory role through inhibition of the hemolytic capacity of C1q, a human complement protein (22). The relative paucity of functional information on *N. americanus* proteins can be attributed, at least in part, to the difficulty in obtaining parasite material and the absence until 2014 of the published *N. americanus* draft genome. Analysis of the 244 Mb draft genome and the predicted 19,151 genes (11), provided important information about the molecular mechanisms and pathways by which *N. americanus* interacts with its human host. In agreement with published transcriptomes of *N. americanus* and genomes/transcriptomes of other hookworm species, a select number of protein families were over-represented, including SCP/TAPS proteins and different mechanistic classes of proteases with various functions including hemoglobinolysis, and tissue penetration (23–28). Of the >19,000 predicted genes reported in the draft *N. americanus* genome, 8,176 genes had no known InterPro domain. Additionally, more than half of the total proteins had either no blast homology to any gene from the NCBI database (10,771 proteins) or shared identity with a ‘hypothetical’ protein (3,043 proteins). These results highlight the importance of further annotation and refinement of the *N. americanus* genome (29).

In general, the practicality of genomic sequence data is dependent on the accuracy of gene annotation as well as the availability of functional, expression and localization information (30). The high-throughput methods used when annotating a genome are prone to errors, therefore, to validate the predicted protein-coding genes, an analysis of the proteome is essential. Mass spectrometry provides useful data that can be used in a proteogenomic approach to improve genome annotation and identify novel peptides containing predicted protein sequences (31). Herein we perform the first proteogenomic analysis of a parasitic helminth, while also significantly improving the genome annotation and comprehensively characterizing the ES proteome of adult *N. americanus*. Our findings provide valuable information on important families of proteins with both known and unknown functions that could be instrumental in host-parasite interactions, including protein families that might be key for parasite survival and protection of the host against excessive immunopathology. Characterization of these proteins will be useful for the identification of vaccine and drug targets and diagnostics for hookworm infection.

## Experimental procedures

### Ethics

Ethical approval for hamster animal experimentation to obtain *N. americanus* adult worms was obtained from the Federal University of Minas Gerais, Brazil (Protocol# 51/2013). Ethical approval for human experimental infection with *N. americanus* and subsequent culturing of L3 was obtained from the James Cook University Human Research Ethics Committee (ID# H5936).

### Parasite material

Adult *N. americanus* were manually isolated from the intestines of experimentally infected golden hamsters (*Mesocricetus auratus*) upon euthanasia. For the isolation and purification of ES products, the worms were washed 3 times in phosphate-buffered saline (PBS) before being cultured overnight in a humidified incubator at 37°C, 5% CO_2_, in RPMI 1640 (100U/ml penicillin, 100 µg/ml streptomycin sulphate, 0.25 µg/ml amphotericin B). The supernatant was collected the following day and debris was removed by centrifuging the concentrated samples at 1,500 g for 3 minutes in a benchtop microfuge. ES products were concentrated with a 3 kDa cut-off Centricon filter membrane (Merck Millipore) and samples were stored at −80°C until use. For somatic adult extracts, data was used from Tang *et al* (11). In brief, whole worms were ground under liquid nitrogen and solubilized using lysis buffer (1.0% (v/v) Triton X-100 in 40 mM Tris, 0.1% (w/v) SDS, pH 7.4). The extract was filtered through 20µm filter before fractionation.

For isolation of *N. americanus* L3, stool samples were collected from infected human volunteers and cultured as follows. Reverse osmosis (RO) water was added to the stool sample until a thick paste was formed. This paste was then distributed onto moistened Filter Paper (VWR, Standard Grade, 110 mm) in Petri dishes (Sarstedt, 150 mm) and placed in a 25°C incubator for 8 days. Following incubation, the edges of each plate were gently rinsed with RO water to obtain clean L3 preparations. Somatic extracts were prepared by adding 100 µl lysis buffer (3 M Urea, 0.2% SDS, 1% Triton X, 50 mM Tris-HCl) to approximately 6,000 larvae before repeated vortexing and sonication (4°C, probe sonicator, pulse setting) to digest the larvae. The extract was passed through a 0.45 µm filter (Millipore) before in-gel fractionation.

### SDS-PAGE fractionation and in-gel trypsin digestion

A total of 30 µg of *N. americanus* adult ES products was buffer exchanged into 50 mM NH_4_HCO_3_, freeze-dried and resuspended in Laemmli buffer. The sample was boiled at 95°C for 5 minutes and electrophoresed on a 12% SDS-PAGE gel for 40 minutes at 150V. The gel was stained with Coomassie Blue and 20 pieces (approximately 1 mm thick) were cut and placed into Eppendorf tubes. For the in-gel digestion, slices were de-stained and freeze-dried before incubating them at 65°C for 1 h in reduction buffer (20 mM dithiothreitol (DTT), Sigma, 50 mM NH_4_HCO_3_). Samples were then alkylated in 50 mM iodoacetamide (IAM, Sigma), 50 mM NH_4_HCO_3_ for 40 minutes at 37°C, washed three times with 25 mM NH_4_HCO_3_ and digested with 20 µg/ml of trypsin (Sigma) by incubating them for 16 h at 37°C with gentle agitation. Digestion was stopped and peptides were released from the slices by adding 0.1% TFA, 70% acetonitrile. This step was repeated 3 times with pooling of the corresponding supernatants to maximise peptide recovery for each sample. Finally, each sample was desalted with a ZipTip (Merck Millipore) and stored at −80°C until use.

### In-solution trypsin digestion and off-gel fractionation

A total of 70 µg of each *N. americanus* extract (L3 somatic and adult ES products) was buffer exchanged into 50 mM NH_4_HCO_3_ before adding DTT to 20 mM and incubating for 10 minutes at 65°C. Alkylation was carried out by adding IAM to 55 mM and incubating for 45 minutes at room temperature in the dark. Samples were digested with 2 µg of trypsin by incubating for 16 h at 37°C with gentle agitation. Following trypsin digestion, peptides were fractionated using a 3100 OFFGEL Fractionator (Agilent Technologies) according to the manufacturer’s protocol with a 24-well format as described previously (19). In brief, Immobiline DryStrip pH 3-10 (24 cm) gel strips were rehydrated in rehydration buffer in the assembled loading frame. Digested peptides were diluted in dilution buffer to a volume of 3.6 ml and loaded equally across the 24 well cassette. The sample was run at a current of 50 µA until 50 kilovolt hours (kVh) had elapsed. Upon completion of the fractionation, samples were collected, desalted with ZipTip and stored at −80°C until use.

### Mass Spectrometry

The extracts were analyzed by LC-MS/MS on a Shimadzu Prominence Nano HPLC (Japan) coupled to a Triple Tof 5600+ mass spectrometer (SCIEX, Canada) equipped with a nano electrospray ion source. Fifteen (15) µl of each extract was injected onto a 50 mm x 300 µm C18 trap column (Agilent Technologies, Australia) at 60 µl/min. The samples were de-salted on the trap column for 6 minutes using 0.1% formic acid (aq) at 60 µl/min. The trap column was then placed in-line with the analytical nano HPLC column, a 150 mm x 100 µm 300SBC18, 3.5 µm (Agilent Technologies, Australia) for mass spectrometry analysis. For peptide elution and analysis the nano-HPLC pump was initially held at 2% solvent B for 6 minutes followed by a linear gradient of 2-40% solvent B over 80 minutes at 500 nl/minute flow rate and then a steeper gradient from 40% to 80% solvent B in 10 minutes was applied. Solvent B was held at 80% for 5 minutes for washing the column and returned to 2% solvent B for equilibration prior to the next sample injection. Solvent A consisted of 0.1% formic acid (aq) and solvent B contained 90/10 acetonitrile/ 0.1% formic acid (aq). The ionspray voltage was set to 2200V, declustering potential (DP) 100V, curtain gas flow 25, nebulizer gas 1 (GS1) 12 and interface heater at 150°C. The mass spectrometer acquired 250 ms full scan TOF-MS data followed by 20 by 250 ms full scan product ion data in an Information Dependent Acquisition (IDA) mode. Full scan TOF-MS data was acquired over the mass range 300-1600 and for product ion ms/ms 80-1600. Ions observed in the TOF-MS scan exceeding a threshold of 150 counts and a charge state of +2 to +5 were set to trigger the acquisition of product ion, MS/MS spectra of the resultant 20 most intense ions. The data was acquired using Analyst TF 1.6.1 (SCIEX, Canada).

### Proteogenomics

The mass spectrometry raw data was searched against the *N. americanus* protein database using SequestHT algorithm in Proteome Discoverer 2.1 (Thermo Scientific, Bremen, Germany). Trypsin was used as the protease, allowing a maximum of two missed cleavages. Carbamidomethylation of cysteine was specified as a fixed modification, and oxidation of methionine was included as variable modifications. The minimum peptide length was specified as 6 amino acids. The mass error of parent ions was set to 10 ppm, and for fragment ions it was set to 0.05 Da. Protein inference was based on the rule of parsimony and required one or more unique peptides. False Discovery Rate (FDR) of 1% at peptide and protein levels was applied. The unassigned spectra from the protein database search were further searched against the six-frame translated genome database of *N. americanus* (11). The genome sequences were downloaded from WormBase ParaSite release 9 (http://parasite.wormbase.org/Necator_americanus_prjna72135/Info/Index) in FASTA format and translated in all six frames using in-house python scripts. All the sequences greater than 10 amino acids between any two stop codons were added to the database. The search was again performed using SequestHT with precursor mass tolerance of 50 ppm and fragment ion tolerance of 1 Da. Carbamidomethylation of cysteine was specified as a fixed modification, and oxidation of methionine was included as variable modifications. Results were obtained at 1% FDR for both protein and peptide level.

The identified peptides in the search against the six-frame translated genome were mapped back to the protein database using standalone BLAST. Any sequences that mapped 100% to the protein database were discarded. The filtered peptides were mapped to the *N. americanus* genome using the standalone tblastn program (11). Peptide identifications that unambiguously mapped to a single region in the genome, also known as Genome Search Specific Peptides (GSSPs), were considered to perform proteogenomics-based annotation of novel coding regions. The known gene annotation GFF3 was downloaded from WormBase ParaSite for *N. americanus* (11). Using in-house scripts, we categorized the GSSPs into various categories: intergenic, intronic, exon-extension, alternative frame, N-terminal extensions or repeat regions with respect to the known regions from GFF3. Additionally, for all the intergenic peptides, we determined if there was a potential open reading frame (ORF) within the stretch of amino acids. Each of the spectra was further manually validated.

### Genome annotation

The genome annotation from *N. americanus* (11) was updated using the MAKER pipeline v2.31.8 (32). The genome assembly (GenBank assembly accession: GCA_000507365.1) was softmasked for repetitive elements with RepeatMasker v4.0.6 using a species-specific repeat library created by RepeatModeler v1.0.8, RepBase repeat libraries (33) and a list of known transposable elements provided by MAKER (32). From the NCBI Sequence Read Archive (SRA), *N. americanus* RNA-Seq data (11) (Adult: SRR609895, SRR831085, SRR892200; L3: SRR609894, SRR831091,SRR89220) were obtained. After adapter and quality trimming using Trimmomatic v0.36 (34), RNA-Seq reads were aligned to the genome using HISAT2 v2.0.5 (35) with the --dta option and subsequently assembled using StringTie v1.2.4 (36). The resulting alignment information and transcript assembly were used by BRAKER (37) and MAKER pipelines, respectively, as extrinsic evidence data. In addition, protein sequences from SwissProt UniRef100 (38) and WormBase ParaSite WS258 (39) (*Ancylostoma ceylanicum* PRJNA231479, *Brugia malayi* PRJNA10729, *Caenorhabditis elegans* PRJNA13758, *Onchocerca volvulus* PRJEB513, *Pristionchus pacificus* PRJNA12644, *Trichinella spiralis* PRJNA12603 and *Strongyloides ratti* PRJEB125) were provided to MAKER as protein homology evidence. Following the developer’s recommendation (40), the protein-coding gene models of Tang *et al.* (11) were passed to MAKER as pred_gff to update the models by adding new 3’ and 5’ exons, additional UTRs, and merging split models. This method, however, cannot change internal exons nor create new annotations where evidence suggests a gene but no corresponding model is previously present. To address this shortcoming, additional *ab initio* gene predictions were generated using BRAKER v2.0.1 (37) and passed to MAKER so that the intron-exon model that best matched the evidence could be included in the final annotation set. Within the BRAKER pipeline, the gene prediction tools GeneMark (41) and AUGUSTUS (42) were trained utilizing the *N. americanus* RNA-Seq alignment and protein homology information from *C. elegans*. The GFFPs identified in the present study were used to confirm the validity of proposed annotation changes and resolve competing gene predictions through manual curation. Gene models with no evidence support were not included in the final annotation build to reduce false positives in the existing annotations. However, *ab initio* gene predictions that encoded Pfam domains as detected by InterProScan v5.19 (43) were rescued to enhance overall accuracy by balancing sensitivity and specificity (32, 44). The completeness of the annotated gene set was assessed using BUSCO v3.0 with Eukaryota-specific single copy orthologs (OrthoDB v9) (45).

### Protein Identification

Mascot version 2.5 (Matrix Science), X!Tandem (The Global Proteome Machine Organisation) version Jackhammer and Comet v2014.02 rev.2 were used to analyze data from the mass spectrometer. Searches were carried out against a database comprised of either the updated genome annotation provided with this study, or the *N. americanus* genome (11), both appended to the common repository of adventitious proteins (cRAP; http://www.thegpm.org/crap/) database (to detect potential contamination). The following parameters were used: enzyme, trypsin; variable modifications, oxidation of methionine, carbamidomethylation of cysteine, deamidation of asparagine and glutamine; maximum missed cleavages, 2; precursor ion mass tolerance, 50 ppm; fragment ion tolerance 0.1 Da; charge states, 2+, 3+ 4+. A FDR of 0.1% was applied, and a filter of greater than 2 significant unique sequences was used to further improve the robustness of data. The mass spectrometry data have been deposited in the ProteomeXchange Consortium via the PRIDE partner repository with the dataset identifier PXD010669.

### Bioinformatic Analysis of Proteomic Sequence Data

Gene ontology (GO) annotations were assigned using the program Blast2GO and Pfam analysis was performed using HMMER (46). Pfam domains were detected at the P<0.001 threshold for the HMMER software. Putative signal peptides were predicted with SignalP v4.1 and transmembrane domains with TMHMM v2.0 (47, 48). REViGO, an online tool, was used to summarise and plot GO terms (49). UpSetR was used to group proteins based on whether they had a GO term with one, two or all of the GO categories (biological process, molecular function or cellular process) (50).

### Similarity analysis

A similarity analysis was carried out based on the Parkinson and Blaxter method using an in-house script (26, 51). Given the difficulty of working with *N. americanus* specifically (in terms of accessibility to samples and establishment of life cycle in other hosts), other hookworms are frequently used to model this human parasite. Data from the secreted proteome of adult *N. americanus* (described herein) was compared to the published secreted proteomes from related adult nematode species including *H. polygyrus, A. caninum* and *Nippostrongylus brasiliensis* (15, 19, 52).

### Protein family similarity visualization

*H. polygyrus, A. caninum* and *N. brasiliensis* SCP/TAPS and protease protein sequences were obtained from their respective published secreted proteomes (15, 19, 52). These sequences were aligned with adult *N. americanus* homologous protein sequences using BLAST. Significant sequence alignments were visualized using Circos (53).

### Phylogenetic Analyses

SCP/TAPS proteins were identified from published secretomes of 6 species of parasitic helminths including *A. caninum, Ascaris suum, H. polygyrus, N. brasiliensis, Trichuris muris* and *Toxocara canis.* SCP/TAPS proteins were identified from *N. americanus* ES products using the proteome data generated in this study. Sequences were sorted into single and double SCP/TAPS domain proteins for individual phylogenetic analysis due to distinct differences described previously (54). A list of the proteins and their respective sequences used for this analysis can be found in Supplemental Data 1. A multiple sequence alignment was carried out using the alignment program MUSCLE. Outliers with poor alignment (long unaligned regions) were detected and filtered out using ODseek. PhyML, a phylogeny software, was used for a maximum-likelihood (ML) phylogenetic analyses of SCP/TAPS amino acid sequences. The tree was visualized with The Interactive Tree of Life (iTOF) online phylogeny tool (https://itol.embl.de/) (55).

## Results

### Proteogenomic analysis and genome annotation

Different proteomes from two life stages (infective L3 somatic extract and adult somatic extracts and ES products) of *N. americanus* were analysed by mass spectrometry in a 5600 ABSciex Triple Tof to perform a proteogenomic analysis of the hookworm genome. After excluding peptides that mapped accurately to the existing protein database entry, a total of 218 novel peptides were identified that did not match any protein sequences of *N. americanus* contained in the annotations of Tang *et al* (11). Of these 218 novel peptides, 83 were found exclusively in adult somatic extracts, 50 exclusively in L3, and 67 exclusively in adult ES, while 10 peptides were found in both the adult somatic extract and ES products, and 8 were found in both adult and larval somatic extracts. No common peptides were identified from adult ES products and larval somatic extract. Newly identified peptides could be grouped into the following six categories: (1) peptides mapping to intergenic regions; (2) peptides mapping to introns; (3) peptides mapping to alternative reading frames; (4) peptides extending gene boundaries (exon extensions); (5) peptides mapping to N-terminal extensions; and (6) peptides mapping to repeat regions (not gene) regions (Figure 1). Of the identified peptides, more than half belonged to group 1, 18% to group 2, 15% to group 3 and a small number to groups 4, 5 and 6 (Figure 1). Group 1 peptides were further analyzed for an ORF including whether or not they contained a methionine; see Supplemental data 2. From this data, one-third of group 1 peptides were found to overlap with a viable ORF, with the shortest having just 21 amino acids and the longest 381 amino acids. The translated protein sequences with their respective peptides, six-frame translated genome (SFG) ID and ORF length are provided in Supplemental data 3.

**Figure 1:**
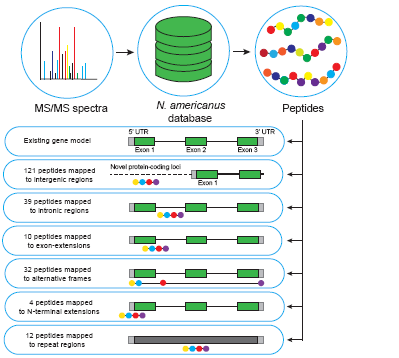
Depiction of the proteogenomic process as well as the types and numbers of peptide corrections identified.

These results highlighted the need for improving the previously published gene models, so we updated the genome annotation of *N. americanus* using a new RNA-Seq based gene calling pipeline as outlined in the Materials and Methods (Table 1). The total number of predicted *N. americanus* genes decreased by 3,425, with substantial increases in both the number of exons and introns. The total coding sequence (CDS) length increased by 2.22 Mb with the mean gene length (including introns and UTRs) nearly doubling from 4.3 kb to 8.1 kb. Furthermore, the percentage of the genome covered by genes increased by 18.3%, and the percentage of detected BUSCOs among the predicted genes increased from 95.7% to 97.4% (with 31% reduction in the number of fragmented BUSCOs). While these improvements are attributable in most part to the use of more sophisticated genome annotation methods utilizing RNA-Seq data and the inclusion of more extensive, up-to-date homologous protein databases, the peptide sequences generated in this study contributed directly to the refinement of 14 gene models. The newly annotated, improved gene models were used in subsequent proteomic analysis of ES products, and are publicly available on Nematode.net (56, 57).

**Table 1:**
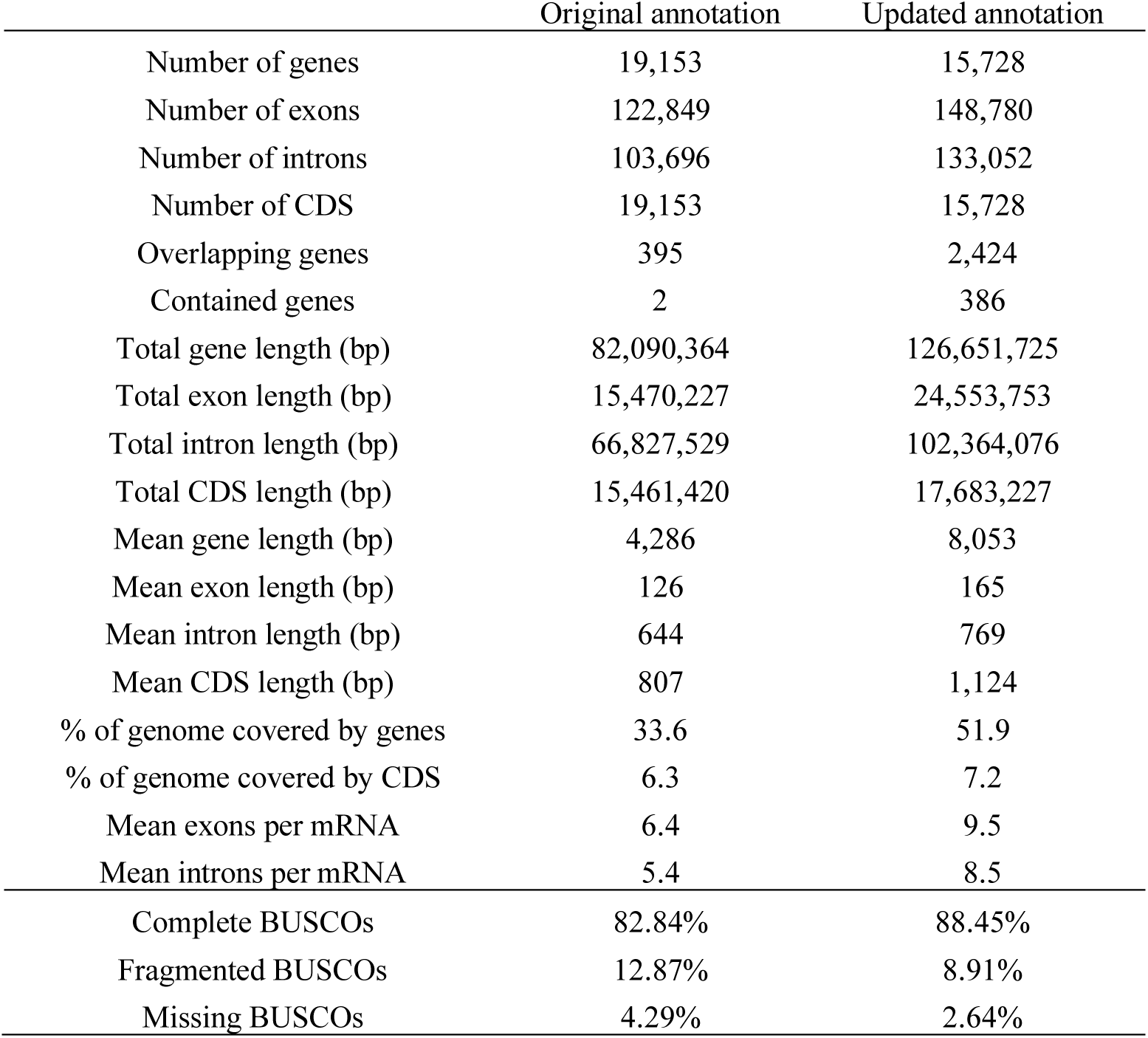
Summary of the updated genome annotation of Necator americanus. CDS - coding DNA sequence.

### Analysis of the ES products from N. americanus adult worms

A comprehensive analysis of the ES products from adult worms was carried out using in-gel and off-gel fractionation and the tryptic peptides were analyzed using LC-MS/MS. Mascot, X!Tandemand Comet searches were carried out against a database including the predicted proteins from the annotated *N. americanus* genome available in this study and the cRAP sequences available at http://www.thegpm.org/crap/. A total of 186 and 141 proteins were identified using Mascot and X!Tandem/Comet, respectively. All ES proteins were identified with at least two unique peptides, at a 99.0% probability and FRD of 0.1%. The two search methods combined found a total of 198 proteins, Supplemental data 4). These 198 proteins were obtained using the updated genome annotation generated as part of this study. In comparison, using the first version of the annotated genome sequence we identified 203 proteins using the same analytical software (11). Using the exponential modified protein abundance index (emPAI) and the newly annotated genome, the most abundant proteins in the ES products of *N. americanus* adult worms were ranked, and the top 30 are shown in Table 2. One emPAI value was listed for each of the digestion methods used (Ingel and Offgel) and their combined value was used to rank the proteins.

**Table 2:**
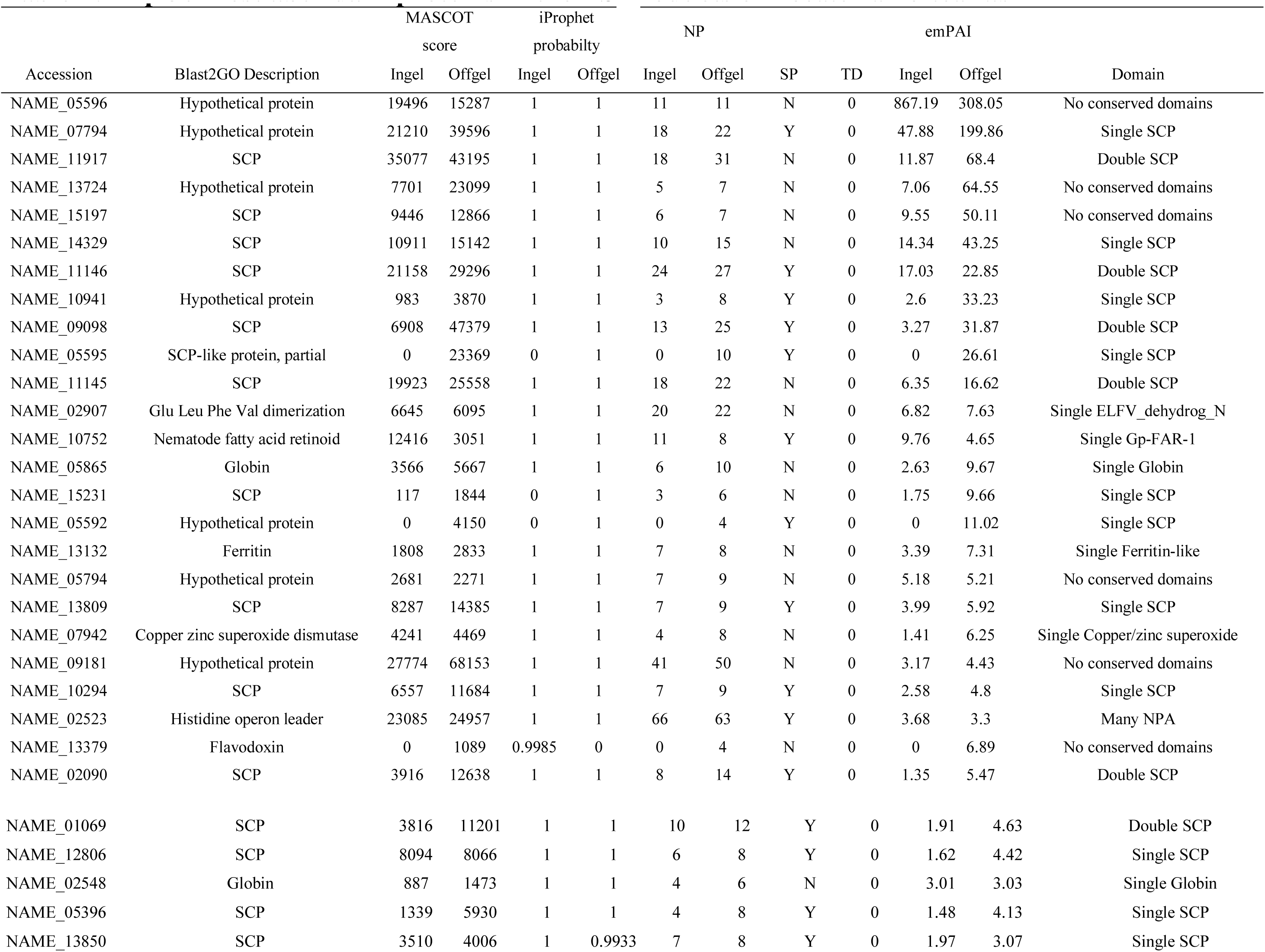
Top 30 most abundant proteins identified with in-gel and OFF-GEL fractionation and ranked using summed exponential modified protein abundance index (emPAI) score. Blast2GO was used to obtain descriptions of each protein. Abbreviations key: NP - number of significant peptides; SP - signal peptide; TD - transmembrane domain; Mascot scores >28 indicate identity or extensive homology (p < 0.05).

The conserved Pfam domains of the 198 ES proteins identified were analyzed. The most abundant protein family in the ES products was the SCP/TAPS family with 54/198 proteins containing a single or double cysteine-rich secretory protein family (CAP) domain (PF00188) (Figure 2B). The top 10 most abundant protein families are displayed in Figure 2B. Many of these proteins were described by Blast2GO as ‘SCP-partial’ or ‘SCP-like’, but for a more standardized annotation in our analysis we have grouped them all as ‘SCP/TAPS’. The second most frequently represented group (with 42 proteins) were proteins with one or more domains of unknown function (DUF). Other abundant families included Ancylostoma secreted protein related (ASPR) proteins and metalloproteases with 35 and 9 of each identified respectively (Supplemental data 5). Despite some published reports classifying ASPRs as SCP/TAPS proteins, they are a diverse set of secreted cysteine rich proteins based on Pfam annotation and therefore we have grouped them separately(28). Of the 198 identified ES proteins, 51% contained a predicted signal peptide. Supporting the accuracy of the new gene model, 96 of the identified adult ES proteins were predicted to contain a signal peptide compared to just 75 using the previous genome annotation.

**Figure 2:**
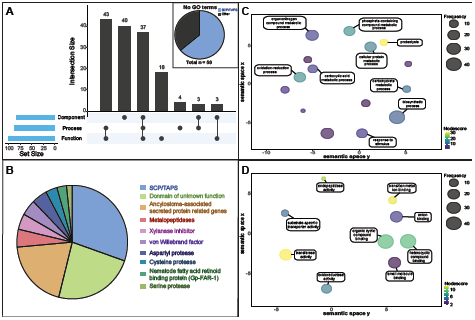
**(A)** UpSetR plot displaying the number of each gene ontology (GO) term categories (biological process, molecular function and/or cellular component) available (Blast2GO) for adult *Necator americanus* excretory/secretory (ES) proteins. Proteins with no available GO terms are broken down in a pie chart into ‘SCP/TAPS’ proteins ‘other’. **(B)** Top 10 most abundant protein families in the ES products of adult *N. americanus*. **(C)** Biological processes of adult *N. americanus* ES proteins ranked by nodescore (Blast2GO) and plotted using REViGO. Semantically similar GO terms plot close together, increasing heatmap score signifies increasing nodescore from Blast2GO, while circle size denotes the frequency of the GO term from the underlying database. **(D)** Molecular functions of adult *N. americanus* ES proteins ranked by nodescore (Blast2GO) plotted using REViGO.

The adult *N. americanus* ES proteins were annotated using Blast2GO (46). In total, 30 GO terms were identified (following the removal of parent child redundancy) belonging to one of the three GO database categories: biological processes, molecular function or cellular component (Figure 2A, for raw data see Supplemental data 6). Blast2GO returned biological process GO terms for 87/198 proteins, molecular function GO terms for 99/198 proteins and cellular component GO terms for 83/198 proteins (Figure 2A). The most prominent biological process term was proteolysis (Figure 2D), with 15% (29 proteins) of total ES products being involved in a proteolytic process. The most prominent single molecular function term was “hydrolase activity” followed by “peptidase activity” and “metal ion binding” (Figure 2C). Interestingly, 50/198 proteins did not return any GO terms (Figure 2A). Of these, SCP/TAPS proteins made up 64% with no known biological process, molecular function, or cellular component. This highlights a significant knowledge gap surrounding SCP/TAPS produced by helminths.

### Similarity analysis of the ES products from different gastrointestinal nematode species

A similarity analysis of ES proteomic data from *N. americanus* and three of the most commonly used animal models for human hookworms, *A. caninum, H. polygyrus* and *N. brasiliensis* was carried out (Figure 3). A total of 15, 10 and 1 *N. americanus* ES proteins had unique homology to ES proteins from *A. caninum, H. polygyrus* and *N. brasiliensis* respectively. This included one SCP/TAPS protein (NAME_13724) which was similar only to an *H. polygyrus* protein, while 3 of the proteins were similar only to *A. caninum* ES proteins with domains of unknown function (NAME_07734, NAME_09181, NAME_09182). Seven proteins shared homology with only *A. caninum* and *H. polygyrus* proteins, 11 shared homology with only *A. caninum* and *N. brasiliensis* proteins and 7 shared homology with only *H. polygyrus* and *N. brasiliensis* proteins. One hundred and twenty-two *N. americanus* proteins shared different degrees of homology with proteins from

**Figure 3:**
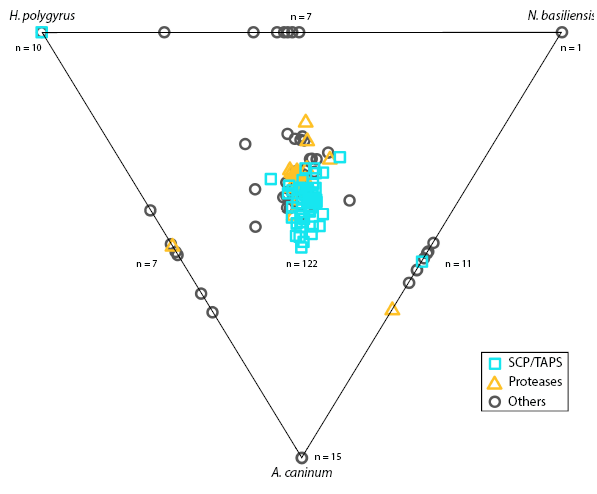
ES products from *Ancylostoma caninum, Heligmosomoides polygyrus, and Nippostrongylus brasiliensis* were compared with the excretory/secretory proteome of *Necator americanus* and displayed in a Simitri plot. SCP/TAPS are represented by an orange square, proteases by a teal-colored triangle and any other protein by a grey circle. Points on the diagram triangle represent sequences which only had similarity to the labelled species. Points along the edges of the triangle are sequences which had similarity to two of the three species (given at the respective ends of the edge). Any sequence in the middle area of the triangle represents a sequence with similarity to all three compared species.

### A. caninum, H. polygyrus and N. brasiliensis

From the 54 SCP/TAPS proteins found in *N. americanus* ES products, one (NAME_13724) shared homology with a protein found in only *H. polygyrus*, while another SCP/TAPS protein (NAME_15177) shared homology with proteins from both *A. caninum* and *N. brasiliensis*. Fifty-one (51) of 54 SCP/TAPS proteins were similar to all compared species, leaving a single SCP/TAPS protein (NAME_11218) which did not have homology to any SCP/TAPS protein from the compared species. Twenty-five *N. americanus* ES proteins did not have homology to any proteins in the secretome of *A. caninum, H. polygyrus* or *N. brasiliensis.* Notable proteins among these 25 were NAME_01848 (aspartyl protease) and NAME_05081 (zinc metalloprotease). Of the 26 proteases in the *N. americanus* ES products, 22 had homologs in all compared species, while one serine protease (NAME_06735) only had homologs in *H. polygyrus* and *A. caninum* and one metalloprotease (NAME_00535, peptidase family M1) only had homologs in the ES products of *A. caninum*, and *N. brasiliensis*. One zinc metalloprotease (NAME_05081) and one serine protease (NAME_01250) were only found in the ES products of *N. americanus* and therefore did not have any similarity to ES products from the other species.

### Homology analysis of SCP/TAPS and proteases in the ES products of N. americanus

Since SCP/TAPS proteins and proteases from *N. americanus* numerically dominate the ES protein dataset and likely play key roles in infection, migration and parasite establishment, we performed an in-depth analysis of these families of proteins between the human and three model gastrointestinal nematode species. Adult *N. americanus* ES SCP/TAPS protein sequences were aligned with homologs from *H. polygyrus, A. caninum* and *N. brasiliensis* ES SCP/TAPS protein sequences using BLAST. Sequences which aligned with maximum scores >36 were visualized using Circos (Figure 4A). SCP/TAPS from *N. americanus* are more similar to *A. caninum* than *H. polygyrus* or *N. brasiliensis*, as denoted by thicker, darker ribbons (Figure 4A). The sequences, their homologs and the corresponding blast scores are detailed in Supplemental data 7. As with the similarity analysis, NAME_11218 had no significant alignment to any of the compared species. A similar analysis was performed for the proteases from each of the aforementioned species (Figure 4B). These proteases have been grouped together into their respective mechanistic classes: aspartyl (ASP), cysteine (CYS), metallo (MET) or serine (SER) proteases. In general, all three comparator species had high protease sequence homology to metalloproteases from *N. americanus*. Aspartyl protease sequences were more similar between *H. polygyrus, N. brasiliensis* and *N. americanus* than sequences from *A. caninum*. It is interesting to note that *N. americanus* contained more aspartyl proteases than the other nematode species analyzed (*N. americanus* – 8; *A. caninum* – 4; *H. polygyrus* – 6; *N. brasiliensis* – 6). Serine proteases had the lowest sequence homology across all of the compared species and protease subclasses.

**Figure 4:**
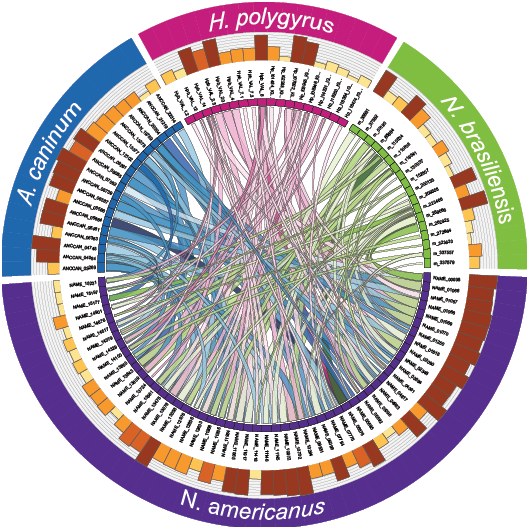
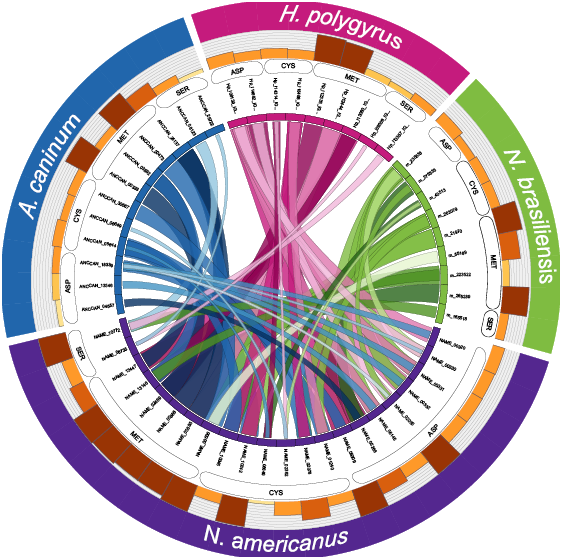
SCP/TAPS proteins in the excretory/secretory products of *Necator americanus* are most closely related to SCP/TAPS proteins in the ES of *Ancylostoma caninum*. SCP/TAPS **(A)** and protease **(B)** protein names are displayed in a circle with *N. americanus* (purple), *A. caninum* (blue), *Heligmosomoides polygyrus* (pink), and *Nippostrongylus brasiliensis* (green). Ribbon thickness is relative to the maximum score obtained in the BLAST search while darker ribbons denote higher sequence percent identity. The corresponding bars provide relative sequence length of each protein. Respective protease mechanistic classes: aspartic (ASP), cysteine (CYS), metallo (MET) or serine (SER).

### Phylogenetic analysis of SCP/TAPS proteins

SCP/TAPS sequences of 6 parasitic nematodes were obtained from published secretomes and compared to *N. americanus* SCP/TAPS identified in this study. Sequences were grouped by whether they had a single or double domain and then aligned using MUSCLE and PhyML. From the 7 total species, 232 SCP/TAPS were reported with 134 single-domain proteins and 98 double-domain. For the single SCP/TAPS-domain proteins an unrooted tree was generated. The analysis identified seven main clades. Three of these clades consisted entirely of sequences from *N. americanus* and *A. caninum*, while sub-clades in 3 of the other main clades followed a similar trend. A majority of *T. muris* single-domain sequences formed a sub-clade with *A. suum* and *T. canis*, grouping together the non-clade V helminths. An unrooted tree was also generated for double domain sequences. The analysis identified five main clades. *N. americanus* SCP/TAPS clustered almost exclusively with sequences from *A. caninum* again, representing one main clade and several sub-clades. *H. polygyrus* and *N. brasiliensis* also formed a number of distinct sub-clades. *T. canis*, the only non-clade V helminth, was reported to only produce one double-domain SCP/TAPS protein in its ES products which was not closely related to any of the compared sequences. Interestingly, no double-domain SCP/TAPS were reported for *A. suum* or *T. muris*. Another trend across both single and double domain trees was the diversity of evolution between SCP/TAPS within a single species. For example, the single-domain tree included a sub-clade of 7 *N. americanus*-only sequences while other *N. americanus* sequences had more similarity to mouse hookworm sequences.

## Discussion

*N. americanus* affects more than 400 million people worldwide and is the most important soil transmitted helminth in terms of morbidity (2). The genome of *N. americanus* was sequenced in 2014, providing an important dataset to facilitate efforts to combat hookworm disease; however, the tools available at that time for annotating genes from parasitic helminths were limited. For instance, the number of proteins with a top hit to a ‘hypothetical protein’ present in the original genome annotation was 3,043, corresponding to 15.8% of the total predicted proteins (11). In addition, inferring gene and protein functions for parasitic nematodes is a major challenge as most species are genetically intractable and databases and algorithms are biased towards (free-living) model nematodes (29). Proteogenomics is a relatively new approach in which proteomic data is used to improve genome annotation (31, 58), and although it had never been applied to parasitic helminths (until now), its potential utility in this area has been suggested (59). High-throughput sequencing and gene prediction tools are prone to false-negative and false-positive predictions which can lead to missed genes, false exons or exon boundaries and/or incorrect translational start/stop sites, so knowing the sequences of the proteins expressed by an organism will help to improve gene predictions.

The proteogenomic analysis carried out in this study addresses some of these challenges by improving the characterization of predicted proteins from the annotated *N. americanus* genome. Overall, we identified 121 peptides that map to intergenic regions in the first draft genome sequence for *N. americanus*. Peptides that map to intergenic regions are highly significant as they can lead to identification of novel protein-coding genes or corrections of pre-existing models (31). To investigate whether these peptides are likely to be new genes we checked for any potential ORFs where the peptides map. Thirty-two (32) of the peptides identified mapped to alternative ORFs than those described in the current gene model. While these peptides map to known coding regions, they highlight out-of-frame ORFs which is likely to, once correctly annotated, result in an entirely different protein. Of the total newly identified peptides, 39 of them mapped to introns. Peptides mapping to introns can lead to identification of novel splice isoforms or amendments in gene structure. Peptides mapping to exon extensions and N-terminal extensions were lessabundant, with 10 and 4 peptides respectively. These groups of peptides suggest a possible correction in reading frame or an incorrect start site annotation.

The decrease in gene number seen in this study is in line with other genome annotations. For example, the *Schistosoma mansoni* genome was originally thought to encode 11,809 genes, however further annotation has reduced this number to 10,772 (60). Given that this is the first re-annotation of the *N. americanus* genome since the original draft was published, a substantial decrease in gene number was to be expected, and 15,728 genes is comparable with the predicted gene numbers of other nematode genomes (61). Despite being a parasitic nematode, *N. americanus* has almost 5,000 fewer genes than its free-living relative *C. elegans* (https://parasite.wormbase.org/).

Of particular importance in the characterization of parasite-specific genes is the presence of a signal peptide. Of the secreted ES proteins from *N. americanus*, 51% were predicted to have a signal peptide; this is also in agreement with ES proteomes from other parasitic nematodes (15, 19, 52). The presence of extracellular proteins without predicted signal peptides in the ES products of *N. americanus* could be due to one of three reasons: (a) the protein is secreted via an alternative pathway, including release of parasite exosomes (62–65) or non-classical secretory signals; (b) the lack of full-length RNA transcript sequence to confirm gene model accuracy resulted in an error in the predicted sequence (i.e. truncation or ORF shift); (c) the pre-set D-cutoff threshold of 0.33 in SignalP results in false-negative predictions.

Helminth secretomes represent the molecular host-parasite interface (10), and provide useful insights into the biological strategies employed by these parasites to ensure longevity inside their respective hosts (10). At this interface, ES products have been implicated in numerous roles from initial penetration/invasion and feeding to host immune regulation (10). Obtaining sufficient ES products to generate the proteome described in this study was time consuming due to the difficulty in culturing sufficient quantities of parasite material in hamsters. For this reason, human infection with *N. americanus* is frequently modelled using other hookworms and related nematodes that survive in rodents or larger animals, including *Ancylostoma sp., H. polygyrus* and *N. brasiliensis.* The protein family analysis of *N. americanus* ES products revealed a diverse number of known and unknown domains (391 domains total), which attests to the many biological functions of ES proteins as well as to the lack of information on these proteomes. The ES products of the three comparator species used here had similar protein family profiles with 458 (*A. caninum*), 434 (*H. polygyrus*) and 628 (*N. brasiliensis*) unique domains present. To assess the overall usefulness of these models, a similarity analysis was carried out on their ES products. This analysis relates relative protein sequence similarities in a plot where each of the identified *N. americanus* ES proteins is compared to the ES proteome of the comparator species (19, 51). The majority of the *N. americanus* ES proteins (173/198), including 51/54 SCP/TAPS proteins, had homologs in the ES products of all three comparators, highlighting the relevance and usefulness of all three models. A total of 25 proteins did not have similarity to ES proteins from the other 3 nematodes analyzed. Since *A. caninum, H. polygyrus* and *N. brasiliensis* are animal parasites, these 25 proteins might have evolved to such an extent that they target human-specific pathways. Two unique proteins of interest are the aspartyl protease NAME_01848 (PF00026) and the zinc metalloprotease NAME_05081 (PF01546). Metalloproteases and aspartyl proteases play crucial roles in host tissue penetration and parasite feeding, and as such, proteolytic enzymes from helminths may be of particular interest as vaccine and/or drug targets (66, 67).

The most abundantly represented protein family (54/198; 27%) in *N. americanus* ES products was proteins containing a single or double SCP/TAPS domain (PF00188). This finding aligns with previous work that highlighted the abundance of this protein family in nematode ES products in particular (15, 68). For instance, a total of 45, 90 and 25 SCP/TAPS proteins were found in the ES products from *N. brasiliensis* (45/313; 14%), *A. caninum* (90/315; 29%) and *H. polygyrus* respectively (25/374; 12%). Interestingly, this family of proteins is also abundant in the extracellular vesicles secreted by different nematodes (64, 65). Given that SCP/TAPS proteins from *N. brasiliensis* are almost exclusively secreted by the adult developmental stage (19), these proteins are likely coordinating specific roles in the gastrointestinal tract of the host. In fact, it has also been shown that SCP/TAPS are overrepresented at the transcript level in *N. americanus* adult worms (11). While relatively little functional information is available for SCP/TAPS proteins, neutrophil inhibitory factor (NIF), an SCP/TAPS protein in the ES of *A. caninum*, was reported to abrogate neutrophil adhesion to the endothelium (69). However, a *N. americanus* homolog of this protein was not detected in the current study, despite the presence of a NIF-encoding gene in the draft genome (11). SCP/TAPS proteins are thought to play numerous and diverse roles at the host-parasite interface, from defense mechanisms, normal body formation and lifespan (54). The diverse nature of *N. americanus* SCP/TAPS sequences is evidenced by their phylogenetic relationships (Figure 5A and B). Blast analyses presented in the Circos plot (Figure 4A) reveal varying degrees of sequence homology to SCP/TAPS proteins from *A. caninum, H. polygyrus* or*N. brasiliensis*. These SCP/TAPS proteins should be further explored to understand their roles in *N. americanus*-human host interactions. Given the limited availability of information regarding the function of helminth SCP/TAPS proteins in general, it was unsurprising that GO analyses revealed no known molecular function or biological process for 32/54 of the SCP/TAPS from *N. americanus* ES products, and more studies should be performed to characterize the properties of this intriguing family of proteins.

**Figure 5:**
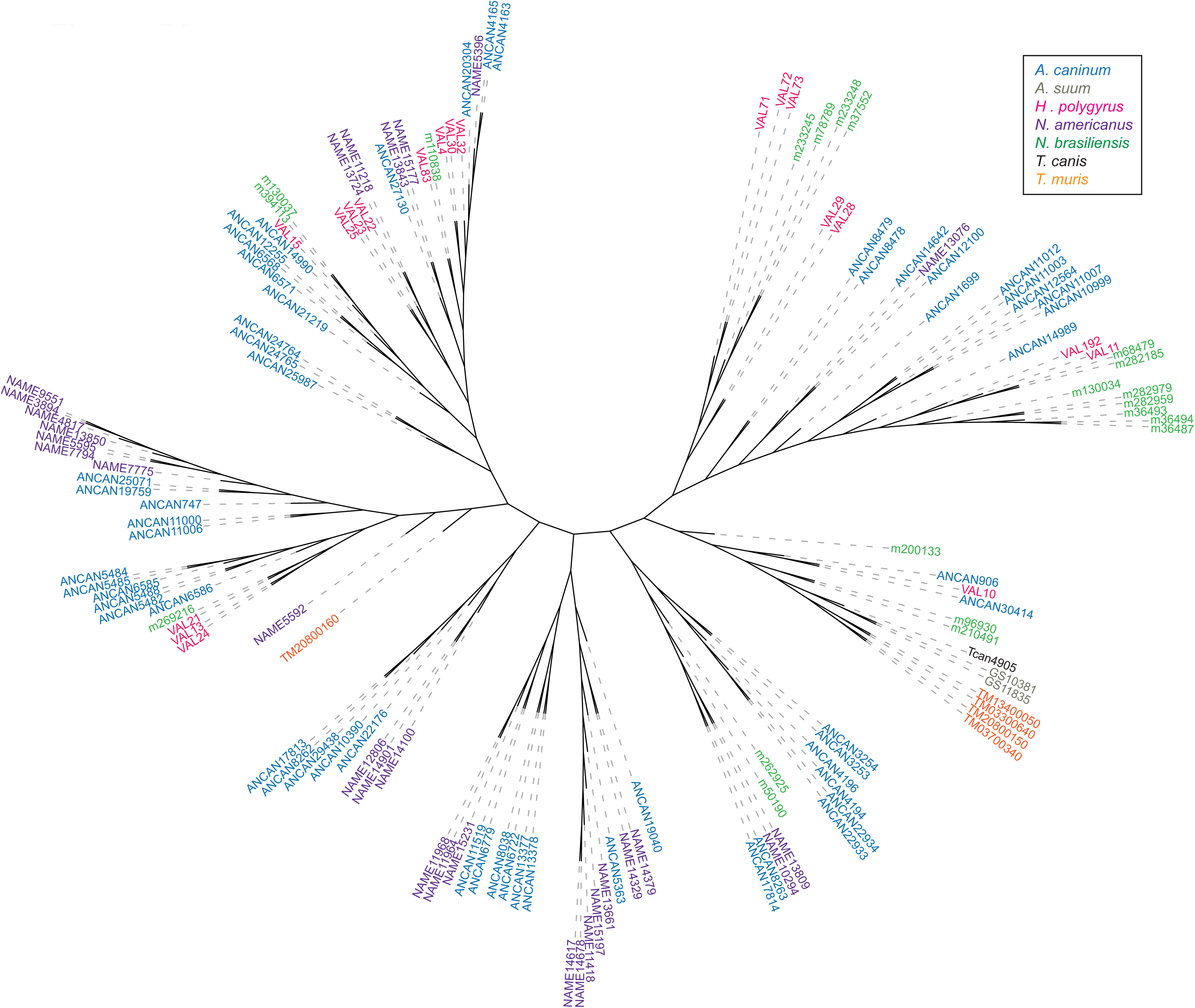
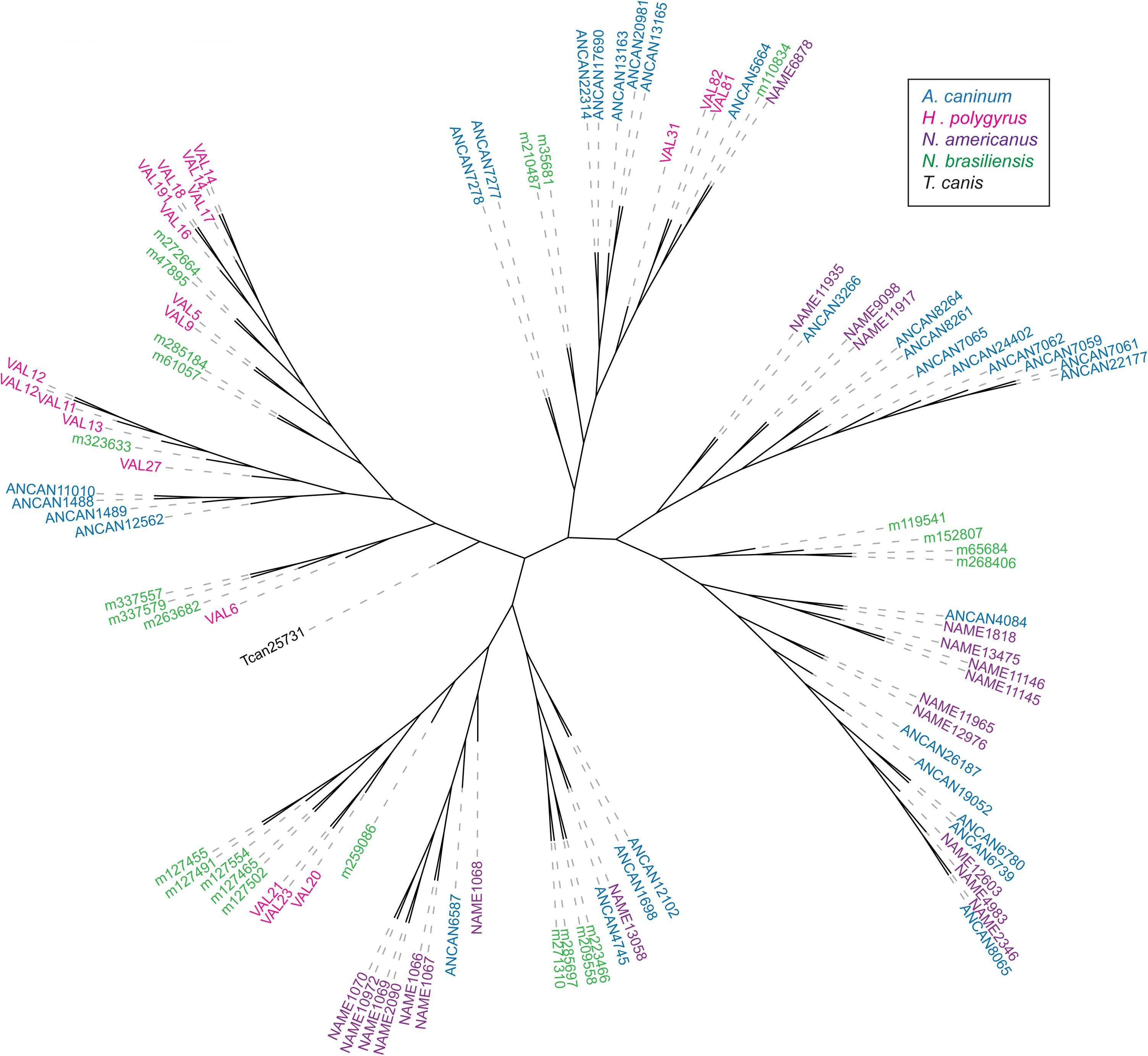
Phylogenetic relationships of **(A)** single-domain and **(B)** double-domain SCP/TAPS proteins determined with MUSCLE alignment software. PhyML was used for a maximum-likelihood phylogenetic analysis and results were visualized with The Interactive Tree of Life (iTOF) online phylogeny tool. *Necator americanus* sequences are highlighted in purple with comparator species each denoted by a different color (see key).

Despite the lack of functional information on the SCP/TAPS proteins in parasitic helminths, of the species we compared in this study, *N. americanus* proteins were generally most similar to those from *A. caninum*, which could simply be a reflection of the phylogenetic similarity between the two species. (Figure 5A and 5B). The trees generated in this study highlight strong clade-specific similarities between SCP/TAPS in the ES products of the compared species. In support of this, the vast majority of the SCP/TAPS proteins came from the four clade V species, while *A. suum, T. muris* and *T. canis* had only 2, 5 and 2 SCP/TAPS proteins respectively. The clustering of *A. caninum* with *N. americanus* and *H. polygyrus* with *N. brasiliensis* supports the notion of host-specific roles for SCP/TAPS. Another trend across both single and double domain trees was the diversity of evolution between SCP/TAPS within a single species. For example, the single-domain tree included a sub-clade of 7 *N. americanus*-only sequences indicating an important human-specific role for this evolutionary cluster. This compares well with a number of other sub-clades which included proteins from the four predominant species. These SCP/TAPS proteins are more likely to share a common function, potentially in host-infection or parasite development.

The phylogenetic analysis strongly attests to the preferred use of *A. caninum* for studying hookworm molecular biology and ES products in general (70). This finding is reinforced by the Circos plot of SCP/TAPS (Figure 4A). Not only was there a greater number of SCP/TAPS homologs in the ES products of *A. caninum* but these proteins also had relatively higher blast scores (denoted by link ribbon thickness) and higher percent identity scores (denoted by ribbon darkness). This type of analysis can prove useful since it also reveals which species to consider for investigating a specific *Necator* SCP/TAPS protein. For example, NAME_13850 has significant sequence homology to a SCP/TAPS protein from each of the compared species; however, the species with the highest sequence homology is *A. caninun* (ANCCAN_19759), making this protein the most relevant to study as a model for NAME_13850. Prior to updating the genome, all three species of nematode used in this comparison had longer average gene sequence lengths than the *N. americanus* SCP/TAPS sequences. In the previous annotation, the average *N. americanus* SCP/TAPS sequence length was 244 predicted amino acids, compared with 355 residues in the updated genome. This was likely due to some of the sequences being truncated, yielding similar results to the published *H. polygyrus* proteome (52).

The Circos plot representing *N. americanus* proteases and homologous proteins from the three comparator species (Figure 4B) provides insight into the high degree of sequence similarity. Unlike the SCP/TAPS Circos analysis, all the *N. americanus* ES proteases had homologs with high similarity in the compared species (average 50% identity between *N. americanus* and comparator species protease). This finding supports the notion of using any one of these three parasites to study *N. americanus* proteases in general. We identified 8 aspartyl, 6 cysteine, 9 metallo and 3 serine proteases in the ES products of *N. americanus*. As the adult stage of *N. americanus* feeds on blood, the high abundance of aspartyl proteases was expected (25, 71). Yet when compared to other species, the human hookworm ES products included more of these proteases. Aspartyl proteases play a fundamental role in the digestion of host hemoglobin and have also been implicated in skin penetration, feeding, and host tissue degradation (72, 73). The finding that *N. americanus* has more of this mechanistic class of proteases than the other nematodes assessed here is likely due to split gene models and/or may be a true gene family expansion. Due to the vital role that aspartyl proteases play in parasite feeding, *Na*-APR-1 an *N. americanus* aspartyl protease, was targeted as a vaccine candidate (74). While we were unable to detect *Na*-APR-1 in the current ES proteome - probably because it is anchored to the gut epithelium (75) - 9 other aspartyl proteases were detected which could potentially be targeted as novel vaccine candidates.

Cysteine proteases, particularly the group belonging to the papain superfamily, are common in nematodes (76). They have been specifically described for their proteolytic activity against hemoglobin, antibodies and fibrinogen in the *N. americanus* lifecycle (77). Similarly, another study highlighted the importance of four cysteine proteases that were upregulated in the *N. americanus* transition from free-living larvae to blood-feeding adult worm, indicating that these proteins are likely to be important for nutrient acquisition (78). Metalloproteases - particularly the astacins - identified in this study are most likely important for larval and adult migration and invasion through human host tissue (79). In support of this, astacin metalloproteases were found to be upregulated in *N. brasiliensis* larvae when compared with adult stage parasites (19). Interestingly, *N. americanus* metalloproteases were reported to inhibit eosinophil recruitment through the cleavage of eotaxin, a potent eosinophil chemoattractant (23). The least abundant family of proteases in the adult *N. americanus* ES products were the serine proteases. Relatively little is known about these proteases from *N. americanus* specifically; however, a serine protease from the whipworm *T. muris* is involved in degradation of the mucus barrier to facilitate feeding (80). Due to the importance of these various proteases in parasite feeding, infection, migration and defense, they represent potential targets for chemotherapies and vaccines to limit infection (81). In the current study, we have provided the first proteogenomic analysis of a helminth parasite, resulting in a more accurate genome annotation. While the above annotations are valuable additions to the current genome, further improvement requires access to substantially more parasitic material including adult stage parasites which are difficult to obtain. Furthermore, we have carried out the first proteomic analysis of the ES products of the human hookworm *N. americanus*. The results presented herein offer significant insight into the validity of these model species while also highlighting differences between these important parasites.

## Acknowledgements

This work was supported by a program grant from the National Health and Medical Research Council (NHMRC) [program grant number 1037304] and a Senior Principal Research fellowship from NHMRC to AL (1117504). The funders had no role in study design, data collection and analysis, decision to publish, or preparation of the manuscript. Work performed at Washington University School of Medicine was supported by NIH-NIAID grant AI081803 to MM. JL was supported by an Australian Postgraduate Award and by the Australian Institute of Tropical Health and Medicine.

## Data availability

The mass spectrometry proteomics data have been deposited to the ProteomeXchange Consortium via the PRIDE partner repository with the dataset identifier PXD010669.

To access the data, please use the below login details.

**Password:** fybW0mgJ

The newly annotated, improved gene models were used in subsequent proteomic analysis of ES products, and are publicly available on Nematode.net.

## Table legends

**Table 1**: Summary of the updated genome annotation of *Necator americanus*. CDS – coding DNA sequence.

**Table 2**: Top 30 most abundant proteins identified with in-gel and OFF-GEL fractionation and ranked using summed exponential modified protein abundance index (emPAI) score. Blast2GO was used to obtain descriptions of each protein. Abbreviations key: NP – number of significant peptides; SP – signal peptide; TD – transmembrane domain; Mascot scores >28 indicate identity or extensive homology (p < 0.05).

## Abbreviations

ASP: Aspartyl
ASPR: Ancylostoma secreted protein related
CAP: Cysteine-rich secretory protein
CDS: Coding sequence
cRAP: Common repository of adventitious proteins
CRISP: Cysteine-rich secretory protein
CYS: Cysteine
DP: Declustering potential
DUF: Domain of unknown function
emPAI: Exponential modified protein abundance index
ES: Excretory/Secretory
FDR: False discovery rate
GO: Gene ontology
GSSPs: Genome search specific peptides
IAM: Iodoacetamide
iTOF: Interactive tree of life
MET: Metallo
ML: Maximum-likelihood
NK: Natural killer
NP: Number of peptides
RO: Reverse osmosis
SCP/TAPS: Sperm-coating protein
SER: Serine
SFG: Six-frame translated genome
SP: Signal peptide
SRA: Sequence read archive
TD: Transmembrane domain

